# Metabolomics-assisted mechanism analysis of γ-PGA synthesis regulated by PhoP in *B. licheniformis*

**DOI:** 10.1101/2023.08.09.552608

**Authors:** Qing Zhang, Shisi He, Wanying Zhu, Fan Yang, Yaozhong Chen, Dongbo Cai, Shouwen Chen

**Affiliations:** State Key Laboratory of Biocatalysis and Enzyme Engineering, Environmental Microbial Technology Center of Hubei Province, College of Life Sciences, Hubei University, Wuhan, 430062, PR China

**Keywords:** Transcription factor PhoP, Poly-γ-glutamic acid, *Bacillus licheniformis*, Metabolomics, GltT, TnrA

## Abstract

Poly γ-glutamic acid (γ-PGA) is a widely used biopolymer whose synthesis relies on external nitrogen sources. PhoP is a global transcription factor that has been reported to be involved in regulation of phosphorus and nitrogen metabolisms, whether PhoP regulates γ-PGA synthesis is worthy of further study. In this study, γ-PGA yield was decreased by 19.4% in *phoP* deletion strain, while PhoP overexpression benefited γ-PGA synthesis in *Bacillus licheniformis*, and the results of transcriptional level, electrophoretic mobility shift (EMSA) and GFP expression assays confirmed the direct positive regulation on γ-PGA synthetase gene *pgsB* by PhoP. Furthermore, based on metabolomic and physiological analysis, we dissected three aspects that γ-PGA synthesis indirectly regulated by PhoP. (i) PhoP influences glutamate transport through positively regulating glutamate transporter GltT. (ii) PhoP influences nitrogen source utilization through negatively regulating nitrogen metabolic repressor TnrA and positively regulating GlnR. (iii) PhoP influences ammonia assimilation through GlnR and TnrA. Together, our study improved metabolic regulatory network of γ-PGA synthesis, and laid a foundation for PhoP regulation nitrogen metabolic network in *Bacillus*.

## 1 Background

Poly-γ-glutamic acid (γ-PGA) is a natural biopolymer formed by the condensation of α-amino and γ-carboxylic acid of glutamic acid(Zhang et al. 2022). The existence of a large number of free carboxyl and carbonyl groups gives it good water solubility. Because of its good biodegradability, nontoxicity, high metal-binding capacity, γ-PGA is widely used in the areas of food, medicine, agriculture, cosmetic and wastewater treatment(Li et al. 2022). γ-PGA is mainly produced by *Bacillus sp.*, and strategies proposed to improve γ-PGA production include strengthening precursor supply, enhancing γ-PGA synthetase expression, blocking by-products synthesis, energy and cofactor regeneration engineering, and transcription factor engineering, etc(Li et al. 2022, Zhang et al. 2022).

Transcription factor can change a wide range of gene expression and products synthesis(Deng et al. 2022, Freyre-González et al. 2013, Zeng et al. 2013). The reported regulatory factors affecting γ-PGA synthesis are mainly related to environmental changes, including ComP-ComA(Kobayashi et al. 2001), DegS-DegU(Hu et al. 2022b), DegQ(Do et al. 2011, Hong et al. 2019), and SwrA. Tran et al. found that inactivation of gene *comP* led to a defect in γ-PGA production in *B. subtilis* 168(Tran et al. 2000). Phosphorylation of ComP activates ComA, which further activates DegQ for γ-PGA synthesis. Besides, DegU~Pi can bind to the −35 region of *pgs* through EMSA assay(Ohsawa et al. 2009), which confirmed its direct regulation on γ-PGA synthesis. Based on the quorum-sensing system, a growth-coupled PhrQ-RapQ-DegU system was constructed, which could dynamically regulate γ-PGA synthesis in *B. subtilis* 168, and finally increased γ-PGA synthesis rate by 6.53 times in a 3 L reactor(Hu et al. 2022a).

PhoR-PhoP is an important two-component regulatory system related to phosphorus metabolism in *Bacillus sp*(Aggarwal et al. 2017, Allenby et al. 2005, Santos-Beneit 2015). For *B. subtilis*, transcription factor PhoP could regulate phosphorus metabolism genes *phoB*(Abdel-Fattah et al. 2005, Liu et al. 1997), *phoA*^18^ and *pstS*(W Liu et al. 1998, Y Qi et al. 1997), and positively regulates respiratory chain related genes *cytB* (Eldakak et al. 2007), *ctaA*(Schau et al. 2004) and *resD*, cell wall and membrane formation related genes *tuaA* (Liu et al. 1998) and *tagAB*(Allenby et al. 2005), etc. In another well-studied host, *Streptomyces*, PhoP was reported to participate in both primary and secondary metabolisms as a global regulator(Salzberg et al. 2015, Santos-Beneit 2015), which can negatively regulate RNA polymerase ω factor gene *rpoZ*(Santos-Beneit et al. 2011), cell respiration and differentiation, antibiotic biosynthesis related genes(Martin et al. 2017). It is worthing noting that PhoP could also affect the expression levels of nitrogen metabolism genes *glnR*, *glnA*, *glnII* and *amtB-glnK-glnD* operon(Martin et al. 2011). Among them, the combined modification of GlnR and PhoP could comprehensively regulate nitrogen and phosphorus metabolisms in *S. coelicolor*, and participated in the efficient biosynthesis of multiple intermediate and secondary metabolites, such as S-adenosyl-methionine(Pei et al. 2022) and erythromycin A(Pei et al. 2022, Xu et al. 2019). The γ-PGA synthesis in *B. licheniformis* is heavily depending on external nitrogen source, whether PhoP regulates γ-PGA synthesis in *Bacillus* has not been reported.

In the last 30 years, there has been an increasing number of studies on γ-PGA synthesis using *Bacillus* as the host bacteria(Cai et al. 2018, Zhang et al. 2023), however, the undissected metabolic regulatory network of γ-PGA seriously hinders engineering breeding for γ-PGA production. In this study, we found that PhoP plays an important role in γ-PGA synthesis in *B. licheniformis*, and further dissected regulated influenced aspects using metabolomics and physiological analyses. This study obtained γ-PGA high-yield strain through transcription factor engineering, and laid a foundation for analyzing the regulation network of γ-PGA synthesis in *Bacillus*.

## 2 Results

### 2.1 Effects of *phoP* overexpression and deletion on γ-PGA production

In this study, transcription factor PhoP was overexpressed in *B. licheniformis* WX-02 to obtain strain WX-02/pHY-phoP. After 30 h cultivation in γ-PGA medium, compared with control strain WX-02/pHY300 (21.82 g/g), γ-PGA yield produced by WX-02/pHY-phoP (27.65 g/g) was increased by 26.7% (*P*<0.005) (**Fig 1A**), suggested that PhoP overexpression benefited γ-PGA production. Furthermore, gene *phoP* was deleted in the original strain WX-02, and γ-PGA yield of resultant strain WX-02ΔphoP was 17.86 g/g, decreased by 19.4% compared to WX-02 (*P*<0.005). In addition, the decrease of γ-PGA yield was alleviated after the complementation of *phoP*, γ-PGA yield of WX-02Δ phoP::phoP returned up to 20.83 g/g, the difference between WX-02 has not statistically significant (*P*>0.05) (**Fig 1A**). Thus, these above results implied that transcription factor PhoP indeed affect γ-PGA biosynthesis.

**Fig 1.**
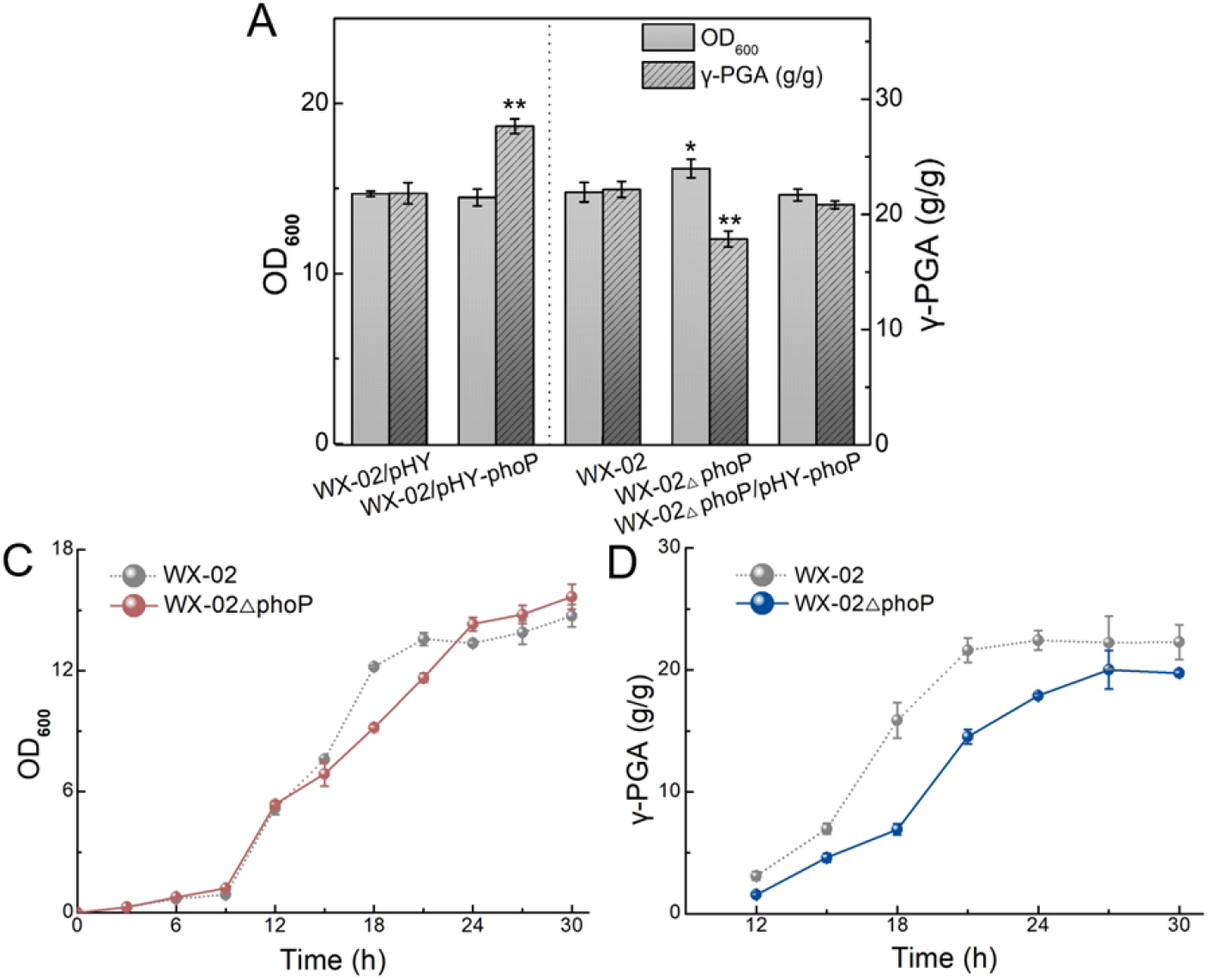
Effects of PhoP overexpression and deletion on γ-PGA production. **A** γ-PGA yields and cell biomass of recombinant strains with overexpression, deletion and complement of gene *phoP* and the controls. **B** Growth curves of *phoP* deficient strain and control strain during γ-PGA fermentation. **C** γ-PGA yield curves of *phoP* deficient strain and its control.

In the further growth curve detection during γ-PGA fermentation, cell biomass of WX-02ΔphoP slowed down in mid-late logarithmic phase (12-18 h), and increased in stable phase (24-30 h) (**Fig 1B**). As for γ-PGA yield, the largest difference was observed around 18 h, only 6.42 g/g γ-PGA was produced by WX-02ΔphoP, which was 60% lower than WX-02 (16.12 g/g) (**Fig 1C**). Therefore, it can be speculated that transcription factor PhoP may affect γ-PGA biosynthesis varies in multi-aspect.

### 2.2 Positive regulation of γ-PGA synthase gene cluster by PhoP

To understand whether PhoP directly regulates γ-PGA synthesis, we examined the relative transcription level of γ-PGA synthase gene *pgsB* in *phoP* deficient strain WX-02ΔphoP (**Fig EV1**), as well as control strain WX-02 (1.000). Our results found the relative transcription level of *pgsB* was significantly reduced in WX-02ΔphoP (0.574), which was increased in PhoP overexpression strain (1.944) (**Fig 2A**). These results seem positively consisted with the results of γ-PGA yields.

**Fig 2.**
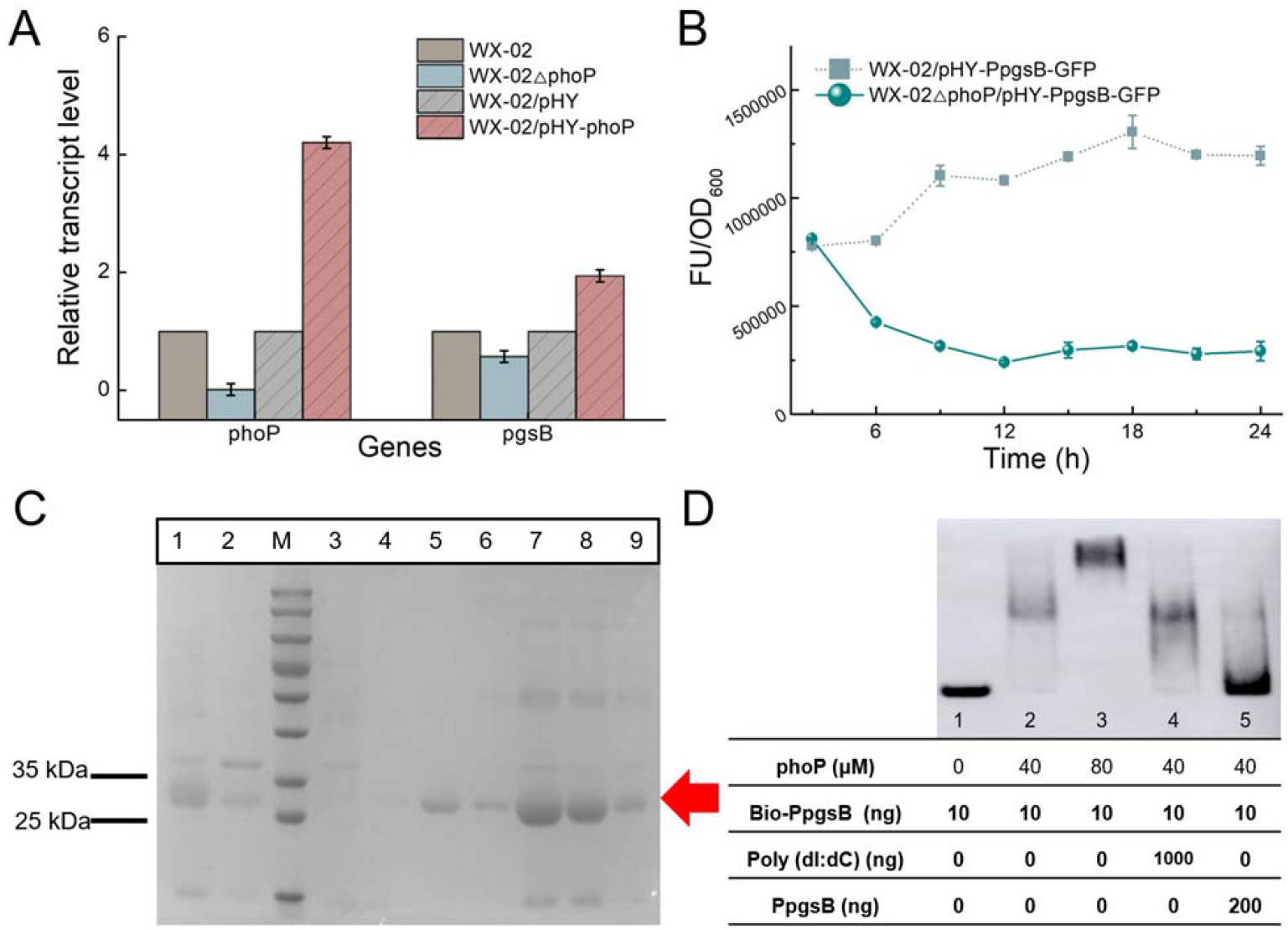
Regulation of γ-PGA synthesis gene *pgsB* mediated by PhoP. **A** RT-qPCR analysis for transcriptional level of γ-PGA synthetase gene *pgsB*. **B** GFP fluorescence assay of *pgsB* promoter in *phoP* deficient strains and its control. GFP was expressed by P_pgsB_ promoter on plasmid pHY300PLK. **C** Detection of PhoP protein purification: Lane 1, samples after IPTG induction, lane 2, samples after cell rupture precipitation, lane 3, flow fluid, lane 4-9, eluent, the monomers size PhoP protein was 27.82 kDa. **D** Electrophoretic mobility shift assays. The target probe is promoter region P_pgsB_ (−183 to +133). Lane 1, negative control, no protein; lane 2, 40 μM PhoP; lane 3, 80 μM PhoP; lane 4, positive control, 40 μM PhoP and 1 μg non-specific competitive DNA poly(dI:dC); lane 5, probe competition reaction, 40 μM PhoP and 200 ng P_pgsB_ without biotin label.

Subsequently, the promoter of gene *pgsB* (P_pgsB_) was used to mediate reporter gene *gfp* expression in WX-02ΔphoP and WX-02, and recombinant strains WX-02ΔphoP/pHY-P_pgsB_-GFP and WX-02/pHY-P_pgsB_-GFP were obtained, respectively. Then, these strains were cultured in LB medium, and samples were taken every 3 hours to detect GFP fluorescence intensities per cell. From 6 h, fluorescence intensity of WX-02ΔphoP/pHY-P_pgsB_-GFP was significantly reduced compared with control (**Fig 2B**), indicated that PhoP positively regulates *pgsB* gene *in vivo*. Electrophoretic mobility shift assays (EMSA) can verify whether target protein and DNA molecular *in vitro*. Biotin-labeled probes Bio-P_pgsB_ and purified PhoP protein (**Fig 2C**) were applied in EMSA assays, Bio-P_pgsB_ contains P_pgsB_ promoter region from −183 to +133. We found that the black band of Bio-P_pgsB_ moved up significantly with the addition of 80 μM PhoP (lane 3), which confirmed the interaction between PhoP with P_pgsB_ *in vitro* (**Fig 2D**). Overall, PhoP regulated γ-PGA biosynthesis by directly activating the expression of γ-PGA synthase gene *pgsB*.

### 2.3 Effects of *phoP* deletion on intercellular metabolic pathways

In bacteria, a wide range of genes were reported to be regulated by regulator PhoP(Han et al. 2023, Martin et al. 2017). In order to find out other aspects that indirectly affected γ-PGA synthesis by PhoP, metabolomics method was adopted to conduct quantitative analysis of WX-02, aimed to dig out the key pathways and target genes that have greater influence, in a bottom-up manner. WX-02ΔphoP and WX-02 were cultured in ME medium supplemented with 10 g/L NaNO_3_, sampled at the mid-log phase (7.5 h) and total OD_600_ was controlled at 10.0 (**Fig 3A**). Based on our results, 98 metabolites with accurate InChIKey values were detected by GC-MS, among which, 73 metabolites were identified according to KEGG database. After standardization, OPLS-DA (orthogonal-partial least squares projection discriminant analysis) was used to analyze the metabolic profile of WX-02 and WX-02Δ phoP, and the above two experimental groups were significantly differentiated (**Fig EV2**).

**Fig 3.**
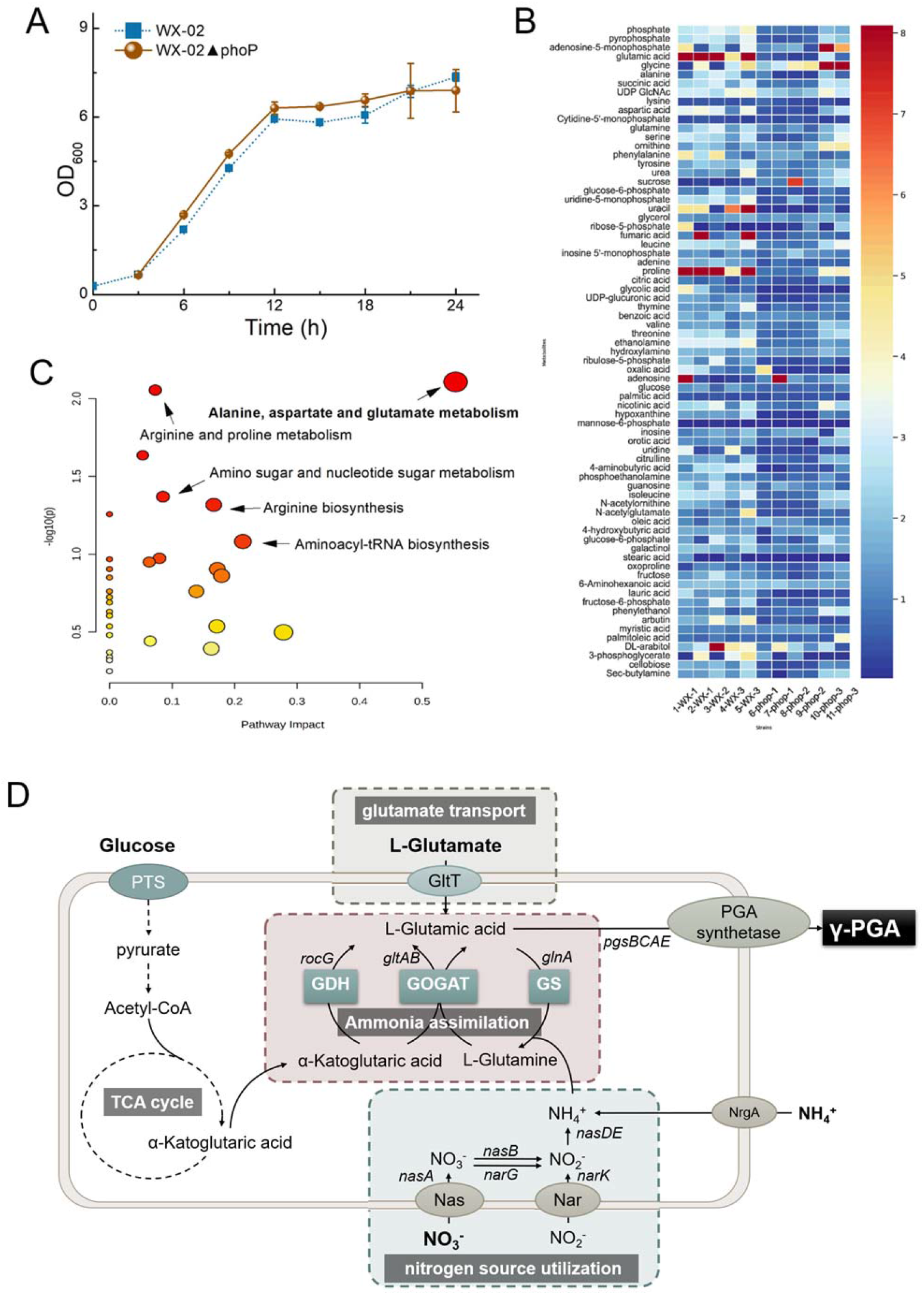
Effects of *phoP* deletion on intercellular metabolic pathways. **A** Growth curve of WX-02ΔphoP and WX-02 in ME medium with additional 10 g/L NaNO_3_. **B** Heat map of differential metabolites. **C** Overview of differential metabolic pathway analysis of WX-02ΔphoP and WX-02. The Y-axis represents the logarithm of P-value, and X-axis represents pathway impact values from pathway topology analysis. **D** Schematic diagram of γ-PGA synthesis in *B. licheniformis*.

PCA analysis using SIMCA analysis software found 33 metabolites with VIP value > 1 (**Fig 3B**) and 19 different metabolites were calculated according to the method described in **2.6** (**Table 1**), among which, phosphate, pyrophosphate, glutamic acid, aspartic acid and glucose-6-phosphate owned the most significant differences. By using MetaboAnalyst software, the above 19 differential metabolites were involved in 35 metabolic pathways of *Bacillus*, and the most influential pathways were alanine, aspartic acid and glutamic acid metabolisms (P = 0.005), followed by arginine and proline metabolisms (P = 0.05), amino sugar and nucleotide sugar metabolisms (P = 0.014), arginine biosynthesis (P = 0.030) and aminoacyl-tRNA biosynthesis (P = 0.030) (**Fig 3C**). The colors (varying from yellow to red) means the metabolite pathways in the data with different levels of significance.

**Table 1.** Different metabolites of WX-02ΔphoP and WX-02.

| **NAME** | **KEGG** | **WX-02 ave** | **▲phop ave** | **p** | **FC** |
| --- | --- | --- | --- | --- | --- |
| phosphate | C00009 | 3.128 | 1.097 | 0.000 | 0.351 |
| pyrophosphate | C00013 | 2.726 | 1.131 | 0.006 | 0.415 |
| glutamic acid | C00025 | 10.409 | 0.870 | 0.002 | 0.084 |
| aspartic acid | C00049 | 2.943 | 1.049 | 0.017 | 0.356 |
| glucose-6-phosphate | C00092 | 1.501 | 0.543 | 0.032 | 0.362 |
| uridine-5-monophosphate | C00105 | 2.665 | 0.802 | 0.001 | 0.301 |
| uracil | C00106 | 5.347 | 0.267 | 0.038 | 0.050 |
| adenine | C00147 | 1.736 | 0.608 | 0.000 | 0.351 |
| proline | C00148 | 10.061 | 1.908 | 0.004 | 0.190 |
| UDP-glucuronic acid | C00167 | 1.378 | 0.422 | 0.000 | 0.306 |
| ethanolamine | C00189 | 3.272 | 0.985 | 0.000 | 0.301 |
| ribulose-5-phosphate | C00199 | 1.793 | 0.440 | 0.034 | 0.246 |
| hypoxanthine | C00262 | 1.651 | 0.417 | 0.004 | 0.252 |
| 4-aminobutyric acid | C00334 | 2.383 | 0.445 | 0.004 | 0.187 |
| fructose | C02336 | 1.716 | 0.855 | 0.032 | 0.498 |
| lauric acid | C02679 | 1.672 | 0.344 | 0.006 | 0.206 |
| fructose-6-phosphate | C05345 | 1.982 | 0.639 | 0.024 | 0.322 |
| arbutin | C06186 | 2.699 | 0.431 | 0.014 | 0.160 |
| cellobiose | C01971 | 1.773 | 0.630 | 0.004 | 0.356 |

As for γ-PGA synthesis in *B. licheniformis*, the precursor glutamic acid can either be transported by extracellular free glutamate or synthesized from glucose metabolism. The former was transferred by glutamate transporter, and the latter is derived from α-ketoglutaric acid of TCA cycle through ammonia assimilation pathway (**Fig 3D**). Thereby, combined with the above biosynthesis pathways, the target metabolic pathways affected by PhoP were focused on the following three aspects, glutamate transport, nitrogen source utilization and ammonia assimilation.

### 2.4 Positive regulation mediated by PhoP on glutamate transport

Glutamate uptake in *Bacillus* is mainly mediated by glutamate/aspartate transporter GltT(Gaillard et al. 1996, Wicke et al. 2019). Then, the mixed D-glutamic acid in the raw material is catalyzed by glutamate racemase to form L conformation, and γ-PGA is synthesized under the action of γ-PGA synthase gene cluster.

We firstly detected the contents of residual glutamate in γ-PGA fermentation process. The residual glutamate of WX-02ΔphoP strains were higher than those of WX-02 (**Fig 4A**). In addition, the glutamate utilization rate of WX-02ΔphoP was slower at the early logarithmic stage (before 18 h), and it was accelerated at late logarithmic stage (after 18 h), suggested that there are different factors affecting the absorption and utilization of glutamate at the early and later logarithmic stage.

**Fig 4.**
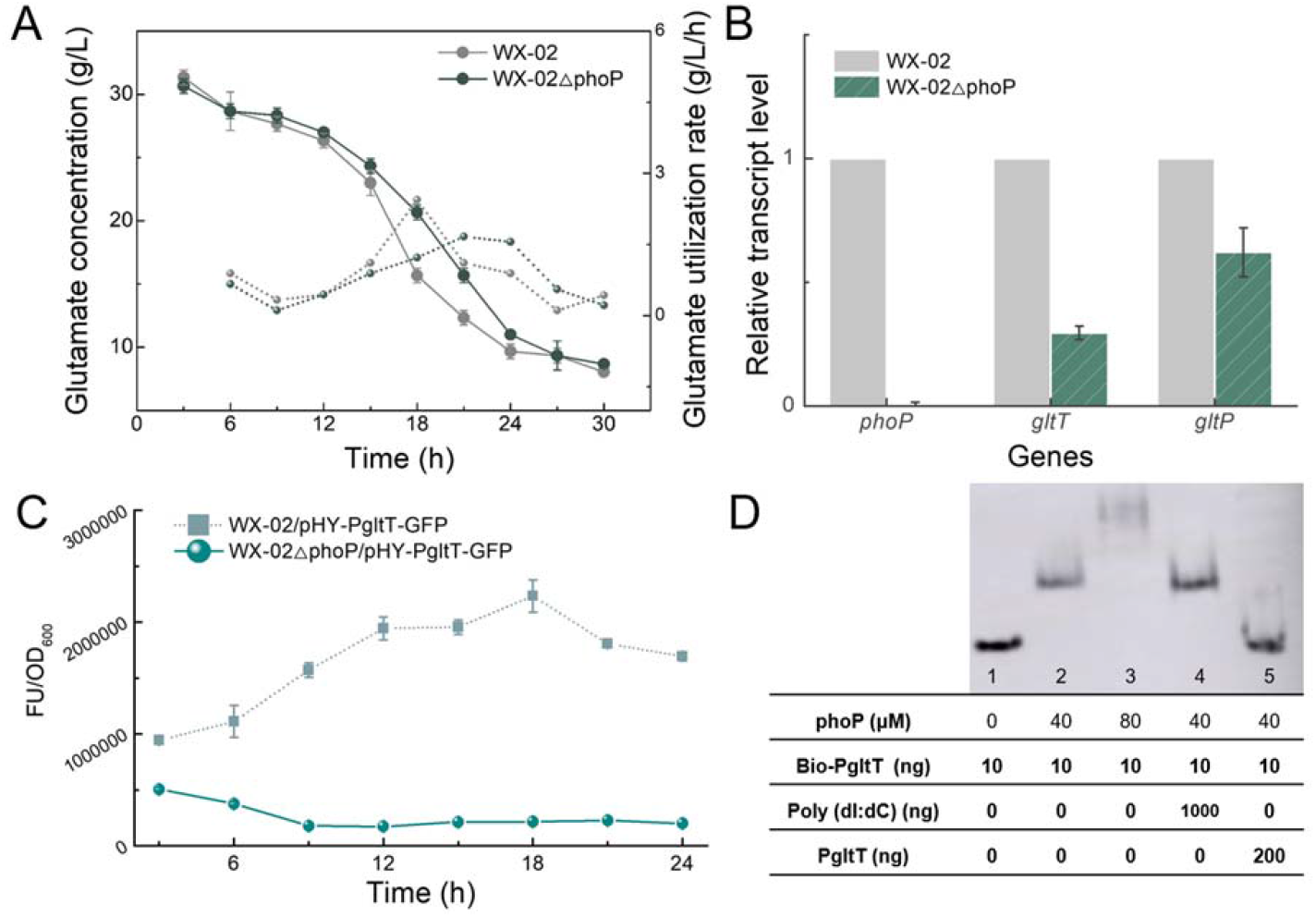
Positive regulation of glutamate symport mediated by PhoP. **A** Line graph of residual sodium glutamate concentration and glutamate utilization rate. **B** RT-qPCR analysis for transcriptional levels of *gltT* and *gltP*. **C** GFP fluorescence assay of *gltT* promoter in *phoP* deficient strains and its control. GFP was expressed by P_gltT_ promoter on plasmid pHY300PLK. **D** Electrophoretic mobility shift assays. The target probe is promoter region P_gltT_(−167 to +254). Lane 1, no protein; lane 2, 40 μM PhoP; lane 3, 80 μM PhoP; lane 4, 40 μM PhoP and 1 μg non-specific competitive DNA poly(dI:dC); lane 4, 40 μM PhoP and 200 ng P_gltT_ without biotin label.

Considering the glutamate transport process, we then investigated the transcription levels of genes *gltT* and *gltP* (**Fig 4B**). Strains WX-02 and WX-02ΔphoP were cultured in γ-PGA production medium, and sampled at 16 h. The relative transcription levels of *gltT* and *gltP* were respectively decreased by 0.027 and 0.621. We assumed that the transcription level of change in greater than double was a significant difference and found *gltT* satisfy this requirement, then we proceeded to examine its intracellular interactions with PhoP proteins. Subsequently, the results of GFP expression (**Fig 4C**) and EMSA assays (**Fig 4D**) confirmed the existence of PhoP protein interaction with *gltT*, suggested that PhoP might affect γ-PGA synthesis through regulating *gltT* expression.

### 2.5 Nitrogen source utilization Regulated by PhoP

The nitrogen sources in γ-PGA fermentation medium include ammonium chloride (NH_4_Cl) and sodium nitrate (NaNO_3_), in addition to sodium glutamate. External ammonium ions entered into cell through transporter NrgAB, and then under the catalysis of GS, they are synthesized to glutamine **(Fig 3D)**. In order to further analyze the influence of PhoP on nitrogen source utilization during γ-PGA production, we first detected the changes of nitrate and nitrite concentration during culture process. Using γ-PGA fermentation medium, the nitrate utilization rate of WX-02ΔphoP (0.220 mg/ml/h) was higher than that of original strain WX-02 (0.157 mg/ml/h) in 3-9 h, but was lower in 12-21 h **(Fig 5A)**. Meanwhile, the rate of nitrite synthesis of WX-02ΔphoP was higher within 3-9 h **(Fig 5B)**, and the large amount of γ-PGA synthesis after 12 h affected biomass detection, so no analysis was performed. Therefore, it is not difficult to speculate that PhoP may inhibit nitrate utilization at the early fermentation stage, and promote nitrate utilization at the middle stage. Therefore, we subsequently sought out some related genes to explore this mechanism.

**Fig 5.**
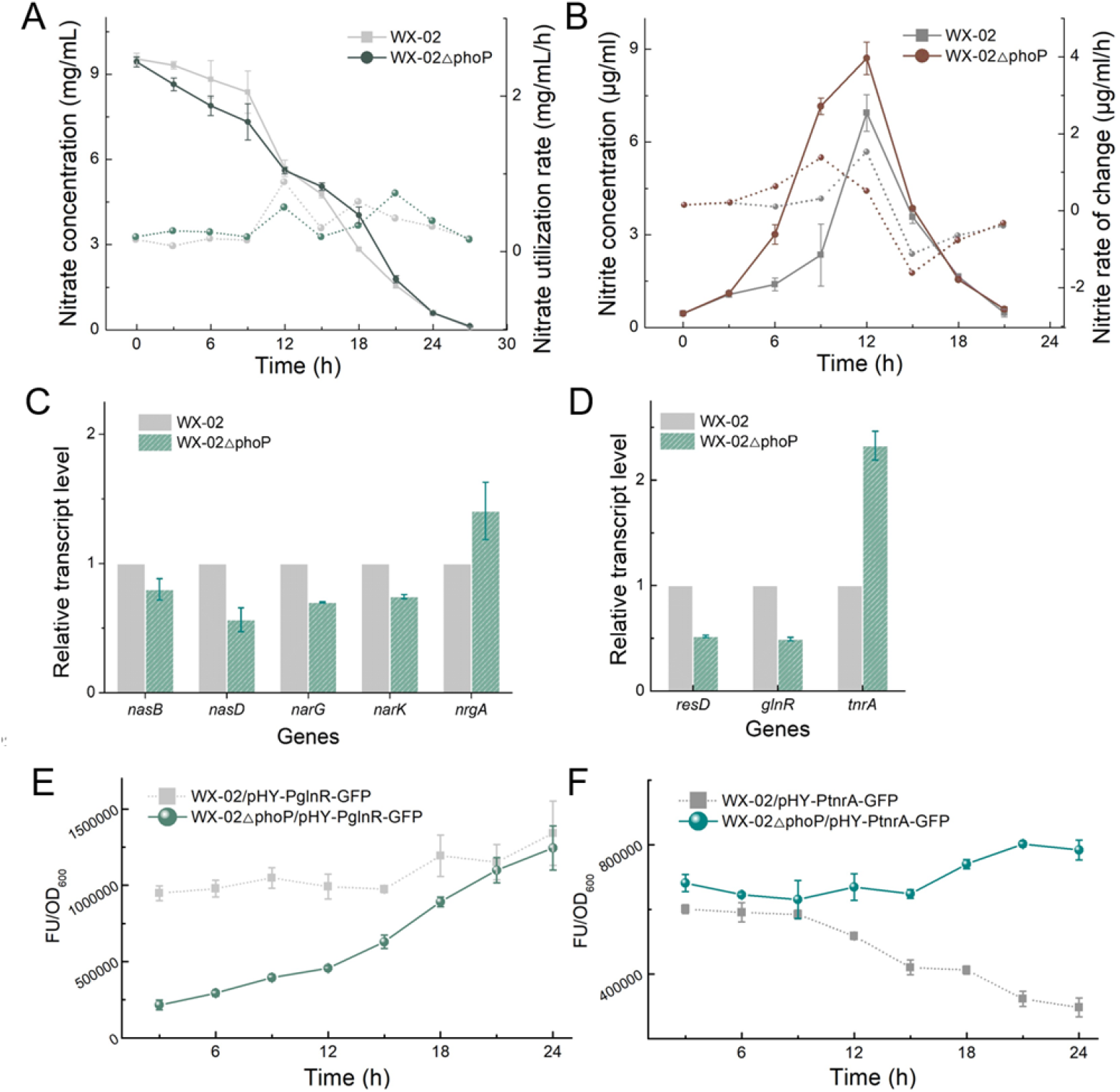
Regulation of nitrogen source utilization mediated by PhoP. **A** Nitrate concentration and nitrate utilization rate in γ-PGA fermentation. **B** Nitrite concentration and nitrite rate of change in γ-PGA fermentation. **C** RT-qPCR analysis for relative transcriptional level of nitrogen source utilization genes involved genes *nasB, nasD, narG, narK* and *nrgA*. **D** RT-qPCR analysis for transcriptional level of nitrogen source utilization genes involve *resD, glnR* and *tnrA*. **E** GFP fluorescence assay of *glnR* promoter activity in *phoP* deficient strains and its control. GFP was expressed by P_glnR_ promoter on plasmid pHY300PLK. **F** GFP fluorescence assay of *tnrA* promoter activity in *phoP* deficient strains and its control. GFP was expressed by P_tnrA_ promoter on plasmid pHY300PLK.

There are two main pathways of nitrate utilization by *B. licheniformis*: nitrate respiration and nitrate assimilation. In nitrate respiration, nitrate is served as the electron acceptor of respiratory chain, which is mainly catalyzed by nitrate/nitrite reductase Nar. While, for nitrate assimilation, nitrate in the medium were transported into cells by assimilate nitrate transporters, and then converted to nitrite by reductase NasBC, and reduced to ammonium ion under the catalysis of NasD, and then participated in the biosynthesis of glutamic acid **(Fig 3D)**. Based on this background, we selectively detected the transcription levels of *nasB*, *nasD*, *narG*, *narK* and *nrgA* in *phoP* deletion strains **(Fig 5C)**. Our results indicated that transcription levels of above genes did not change significantly during γ-PGA fermentation process after *phoP* deletion, exceptionally, the transcription level of gene *nasD* was increased, and further GFP expression experiments also confirmed that *phop*-deficient strain had lower relative fluorescence level mediated by *nasD* promoter **(Fig EV3)**.

It was noted that three transcription factors participated in nitrogen utilization, two-component regulatory system ResD-ResE(Nakano et al. 2000, Nakano et al. 1996), TnrA(Mirouze et al. 2015, Nakano et al. 1998) and GlnR(Zalieckas et al. 2006) regulators. Detection of gene transcription levels in γ-PGA high-yield processes suggests that deletion of *phoP* leads to down-regulation of gene *resD* transcription level **(Fig 5D)**, and this gene has been reported to positively regulate genes *narK* and *narG* expression in nitrate respiration. It is not difficult to deduce that PhoP can promote nitrate respiration by positively regulating *resD* expression during γ-PGA synthesis. Interestingly, *glnR* expression was down-regulated by 50.7%, and *tnrA* expression was up-regulated by 132.6% **(Fig 5D)**, which seemed to indicate that the indirect regulation mediated by regulators GlnR and TnrA had a greater influence on nitrogen source utilization. Whereupon, we further explored the regulatory effect of PhoP on GlnR and TnrA.

In order to further elucidate the relevant mechanism, gene *gfp* was expressed mediated by the promoter of *glnR*(P_glnR_) and *tnrA*(P_tnrA_), respectively, and was transformed into WX-02 and WX-02Δ phoP. Since the produced γ-PGA affects cell turbidity, we used ME medium supplemented with 10 g/L NaNO_3_ for the choice of GFP expression. Samples were taken every 3 hours to detect biomass and fluorescence intensity. GFP fluorescence intensities of WX-02ΔphoP/pHY-P_glnR_-GFP were decreased significantly in the early growth period (0-18 h), and had no significant difference after 21 h (**Fig 5E**). On the contrary, which of WX-02ΔphoP/pHY-P_tnrA_-GFP had no significant difference at the early growth period (0-9 h), but was increased significantly after 12 h (**Fig 5F**). These results confirmed the positive regulation of *glnR* by regulator PhoP, and firstly demonstrated the negative regulation of PhoP on TnrA.

Recently, several researches have discussed the regulation and production application of nitrogen trophic transcription factors GlnR and PhoP, but there are few studies on the relationship between TnrA and PhoP. This conclusion is confirmed again in our subsequent EMSA assays, biotin-labeled *tnrA* probe can produce significant tagging with PhoP protein (**Fig EV4**).

### 2.6 Positive regulation of PhoP on ammonia assimilation through GlnR and TnrA

As the core pathway of nitrogen metabolism, ammonia assimilation includes two pathways, glutamate dehydrogenase (GDH) pathway, glutamine synthesis and glutamate synthase (GS-GOGAT) pathway(Li et al. 2022) (**Fig 3D**). GDH (encoded by *rocG*) in *B. licheniformis* is a NADPH-dependent dehydrogenase that catalyzes the convention of α-ketoglutaric acid from TCA cycle to glutamic acid. GS (encoded by *glnA*) can synthesize glutamine using glutamic acid and ammonia ions, in which ammonia ions can be either transported into cell by transporter NrgAB or generated by nitrogen assimilation. Therefore, GS-GOGAT (encoded by *gltAB*) pathway could achieve ammonia fixation and nitrogen storage. Therefore, WX-02 and WX-02ΔphoP were cultivate in γ-PGA fermentation medium, and transcription levels of ammonia assimilation genes (*rocG, glnA* and *gltA*) were measured at 16 h. Based on results of **Fig 6A**, transcription levels of *rocG, glnA* and *gltA* were decreased by 14.5%, 12.0% and 23.7%, respectively. Ammonia assimilation during γ-PGA production may be positively regulated by PhoP, but this effect seems to be not profound. We noted that in addition to participating in nitrate metabolism, transcription factors GlnR and TnrA are also important regulators in ammonia assimilation. Therefore, we speculated that PhoP indirectly regulated ammonia assimilation through regulators GlnR and TnrA, in which *glnR* expression was down-regulated by 50.7% compared to original strain WX-02, and *tnrA* expression was increased by 132.6% **(Fig 5D)**.

**Fig 6.**
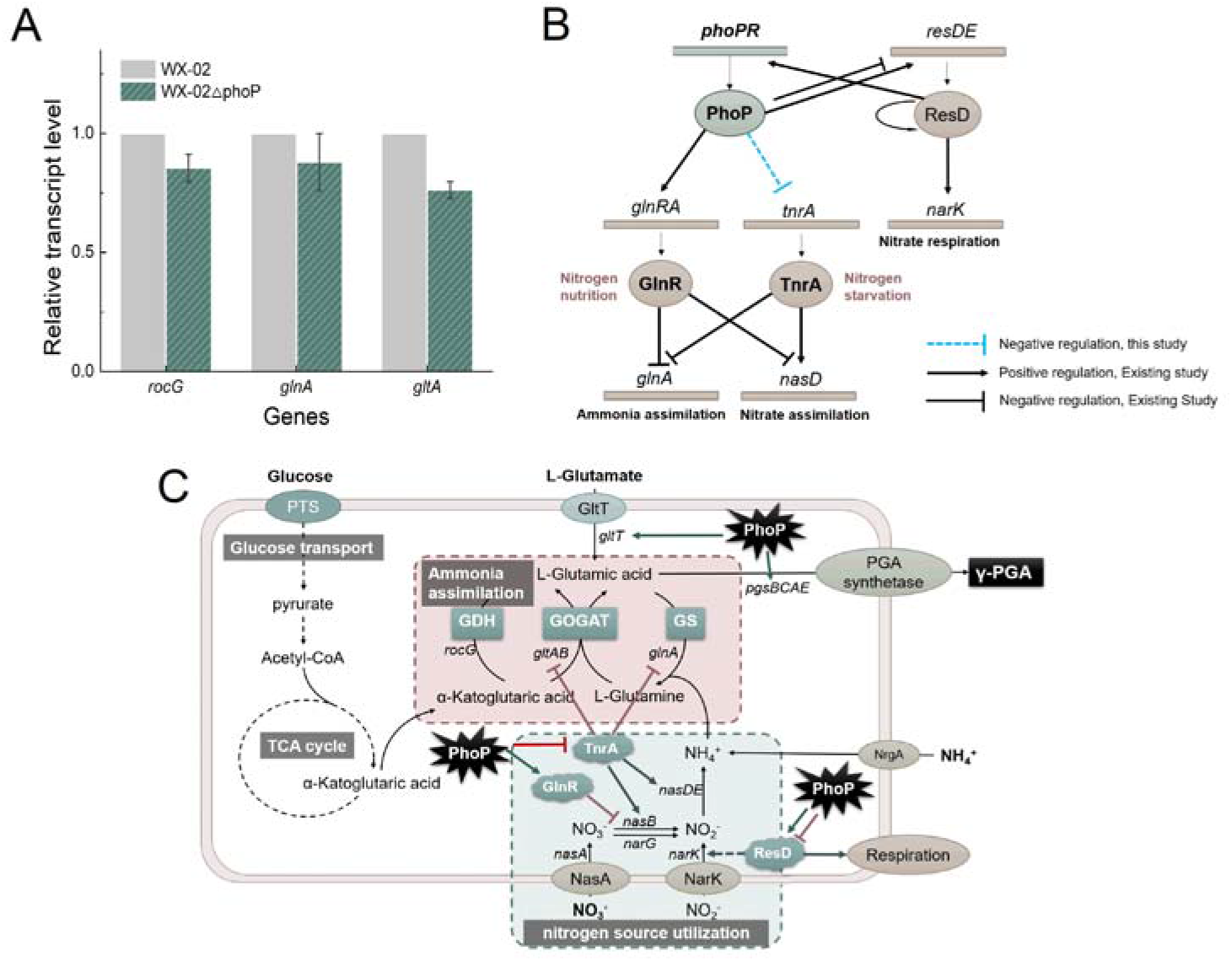
Positive regulation of PhoP on ammonia assimilation through regulators GlnR and TnrA. **A** RT-qPCR analysis for transcriptional level of glucose-phosphotransferase system involve *rocG*, *glnA* and *gltA*. **B** Network model of PhoP comprehensive regulation of nitrogen metabolism. **C** Regulation network diagram of PhoP on γ-PGA biosynthesis.

Combined with the above results and previous researches*(Randazzo et al. 2017)*, we proposed a comprehensive regulation network model of PhoP on nitrogen utilization and ammonia assimilation (**Fig 6B**), and analyzed the regulation network of PhoP on γ-PGA synthesis (**Fig 6C**).

## 3 Discussions

Poly γ-glutamic acid (γ-PGA) is a kind of biological macromolecular with important applications. The precise analysis of its metabolic regulation mechanism is the basis of metabolic engineering breeding to achieve high-level production. However, the lack of research on γ-PGA metabolic regulatory network limited its efficient synthesis. The phosphorous metabolic regulator PhoP has been reported to involving in nitrogen metabolism regulation in *Streptomyces*. Considering that the synthesis of γ-PGA requires a large number of extracellular nitrogen sources, whether PhoP can regulate γ-PGA synthesis was studied in this research. We found that the yield of γ-PGA was increased 26.7% in regulator PhoP overexpression strain, and decreased by 19.4% via deleting regulator *phoP* in WX-02. We further demonstrated that PhoP can directly negative regulate γ-PGA synthase gene *pgsB*, which is the first gene of *pgs* operon. Combined with metabolomic and physiological analysis, we further dissected that PhoP could indirectly regulate γ-PGA synthesis by influencing glutamate transport, ammonia assimilation and glucose transport. Our research laid a foundation for analyzing the regulation network of γ-PGA synthesis in *Bacillus*.

PhoR-PhoP is an important two-component regulatory system related to phosphorus metabolism. In *Streptomyces*, regulator PhoP can activate *glnR*, the gene encoding nitrogen metabolism regulator GlnR. It can be seen that PhoP plays a global regulatory role, and affects the production of a variety of secondary metabolites. Xu et al. proposed a GlnR–PhoP–BldD-*ery* regulatory network to coordinate the primary nutrient metabolism and erythromycin biosynthesis in response to nitrogen/phosphate sources availability in *S. erythraea*(Xu et al. 2019). However, the regulation of regulator PhoP in *Bacillus* mainly focuses on phosphorus metabolism. In the process of exploring the interaction of PhoP on γ-PGA production, this study not only confirmed that PhoP can positively regulate nitrogen nutrition transcription factor GlnR in *B. licheniformis*, but also first discover the negative regulation of PhoP on nitrogen starvation regulator TnrA.

Transcription factors GlnR and TnrA are both nitrogen-responsive, but have opposite regulation effects. GlnR plays a role in nitrogen nutrition, positively regulating glutamine synthase gene *glnA* and negatively regulating assimilatory nitrate reductase gene *nasB*. When nutrient abundance gradually becomes scarce, TnrA began to be transcribed, expressed, inhibited gene *glnR*, and regulated several nitrogen metabolism genes, including negative regulation of glutamate synthase *gltAB* and positive regulation of nitrogen assimilation genes *nasD* and *nasB*. In this study, PhoP can interact with the above two regulators, promoting *glnR* expression at the early stage of γ-PGA fermentation and inhibiting *tnrA* gene expression in the late stage. This temporal regulation indicates that PhoP in *B. licheniformis* can assist in the completion of nitrogen source regulation as a widespread transcription factor.

This study firstly found overexpression of transcription factor PhoP can improve γ-PGA yield of in *B. licheniformis*. PhoP could directly positively regulate γ-PGA synthase gene *pgsB*, and we further analyzed the metabolic mechanism of PhoP indirectly regulating γ-PGA synthesis from glutamate transport, nitrate metabolism and ammonia assimilation. This study improved metabolic regulatory network of γ-PGA synthesis, and provided new ideas for γ-PGA breeding and analysis of PhoP regulatory network.

## 5 Materials and methods

### 5.1 Strains, plasmids and cultivation conditions

All strains and plasmids used in this study are listed in **Appendix Table S1**. *E. coli* DH5α strain was used for vector construction, *E. coli* BL21(DE3) was used for PhoP0 induction expression. *B. licheniformis* WX-02 was used as the original strain for constructing recombinant strains. The plasmid pHY300PLK was applied for constructing PhoP and GFP expression vectors, and T_2_(2)-Ori was used for gene *phoP* deletion, and pET28a was applied for protein induction expression vector construction.

*B. licheniformis* and *E. coli* strains were grown on Luria-Bertani (LB) medium, and corresponding antibiotics (tetracycline 20 μg/mL; kanamycin 20 μg/mL) were added when necessary. The γ-PGA fermentation medium contained 80 g/L glucose, 20 g/L sodium glutamate, 18.4 g/L sodium citrate, 10 g/L NaNO_3_, 8 g/L NH_4_Cl, 1.0 g/L K_2_HPO_4_, 1.0 g/L MgSO_4_·7H_2_O, 1.0 g/L ZnSO_4_·7H_2_O, 0.15 g/L MnSO_4_·7H_2_O, 1.0 g/L CaCl_2_·2H_2_O, pH 7.2. The seeds were cultivated in 250 mL flask containing 50 mL LB for 12 h, then transferred into γ-PGA production medium at a volume ratio of 3%, and cultivated at 230 rpm for 30 h. ME medium contains 20 g/L glucose, 20 g/L sodium glutamate, 12 g/L sodium citrate, 7 g/L NH_4_Cl, 0.5 g/L K_2_HPO_4_, 0.5 g/L MgSO_4_·7H_2_O, 0.04 g/L FeCl_3_·6H_2_O, 0.104 g/L MnSO_4_·H_2_O, 0.15 g/L CaCl_2_·2H_2_O, pH 7.0. All environments were maintained at 37°C except for strain subcultures.

### 5.2 Construction of gene overexpression strain

The method for gene overexpression strain construction was referred to our previously reported research(Wei et al. 2015, Zhang et al. 2023). P43 promoter from *B. subtilis* 168, gene *phoP* and terminator TamyL from *B. licheniformis* WX-02 were amplified. T5 recombination kit was used to connect expression elements and corresponding plasmid vector backbone, and recombinant plasmid pHY-phoP was obtained and electroporated into *B. licheniformis* to obtain PhoP expression strain WX-02/pHY-phoP. Similarly, GFP expression strains were attained by the same method.

### 5.3 Establishment of *phoP* deletion strain

Temperature-sensitive plasmid T_2_(2)-Ori was used for constructing gene *phoP* deletion vector. Firstly, upstream and downstream homology arms of *phoP* were respectively amplified, after Splicing Overlap Extension (SOE)-PCR ligation, and then assembled into T_2_(2)-Ori vector, which step is completed using T5 recombination cloning kit as above. Positive mutants were verified by PCR and DNA sequencing, named as T_2_-ΔphoP. Then, T_2_-ΔphoP was electro-transferred into *B. licheniformis* WX-02, positive clones were cultivated in LB medium with 20 μg·mL^−1^ kanamycin at 45°C, and sub-cultured for three times to obtain the single-crossover recombinants. The recombinants were then grown in LB medium at 37°C with six subcultures, and kanamycin sensitive clones were further confirmed, and the corresponding *phoP* deletion strain was named as WX-02ΔphoP.

### 5.4 PhoP induction expression and purification

The PhoP protein was expressed via pET28a(+) plasmid (Novagen, Denmark). The amplified *phoP* gene was inserted into pET28a using T5 recombination cloning kit, named as pET28a-phoP. Then, pET28a-phoP was transferred into *E. coli* BL21, PhoP protein was expressed after IPTG induction, and purified by Ni-NTA purification kit.

### 5.5 Analytical methods

Cell biomass was attained via determining dry cell weight, and γ-PGA yield was measured by high performance liquid chromatography (HPLC), according to our previously reported method(Zhan et al. 2018). The concentrations of glucose and glutamic acid were detected by SBA-40C bio-analyzer (Academy of science, Shandong, China). The nitrate concentration was detected by salicylic acid method, and nitrite concentration was determined by naphthalamine colorimetric method, as previously reported. For gene transcriptional level analysis, the total RNA was extracted using RNA extraction kit (FORE GENE, China), and first strand cDNA was amplified using RevertAid First Strand cDNA Synthesis Kit (Thermo Fisher, USA). The real-time PCR was performed by using SYBR Select Master Mix (ABI, USA), and *16S rDNA* was used as reference gene to normalize data. The GFP fluorescence intensity were detected by Multi-Mode Microplate Reader (Spectra Max iD3, Molecular Devices, USA), and fluorescence intensity of per cell is the ratio of GFP fluorescence intensity to cell biomass.

### 5.6 Metabolites extraction, GC-MS and metabolomics data analyses

For metabolomics analysis, strains were cultivated in ME medium, and the same biomass was precisely sampled. The quenching, extraction procedure and GC-MS analysis method was carried out according to Yuan et al(Yuan et al. 2019). GC-MS data were first converted into abf format using Reifycs Abf (Analysis Base File) Converter, and MSDIAL software was used to match the above data with Fiehn Bin Base Metabolome Database and output “Height file”^35^. Then, the peak area was normalized with internal standard. The processed data was firstly performed the principal component analysis (PCA) and pair-wise orthogonal partial least squares discriminant analysis (OPLS-DA) by SIMCA 14.1 (Umetrics, Sweden). Metabolite with a VIP value > 1, *p* value < 0.05, and a fold change of > 2 or < 0.5 in each comparison was selected as a differential metabolite. The differential metabolic pathways were calculated using MetaboAnalyst software (http://www.metaboanalyst.ca/MetaboAnalyst/) to analyze the contribution of differential metabolites to Kyoto Encyclopedia of Genes and Genomes (KEGG) pathway. The heat map is produced by HemI (Heatmap Illustrator, http://hemi.biocuckoo.org/). Gene expression level with a fold change of > 2 or < 0.5 was considered as a changed gene.

### 5.7 Electrophoretic mobility shift assay

Biotin probes of target genes are amplified using a 5’ biotin-labeled primer (Sangon Biotech, China). The DNA probe was used for EMSA assays with purified PhoP protein. EMSA assays were completed by chemiluminescence EMSA kits (Beyotime, China). Non-specific competitive DNA is poly(dI:dC) provided by Thermo Fisher. Imaging was done by Amersham Imager 600 (Gelifesciences, USA).

### 5.8 Data analysis

At least three parallels were conducted for each experiment, Origin 8.5 and SPSS 18.0 were used for data processing and analysis.

## Supplementary information

Supplementary data associated with this article can be found in the online version of the paper.

## Acknowledgements

This work has received funding from the National Key Research and Development Program of China (2021YFC2101700), Knowledge Innovation Program of Wuhan-Shuguang Project (2022020801020334), Science and Technology Project of Hubei Tobacco Company (027Y2021-023).

## Author’s contributions

**Q Zhang:** Methodology, Investigation, Data curation, Software, Writing - original draft. **S He:** Methodology, Investigation, Data curation. **W Zhu:** Investigation, Data curation. **F Yang**: Methodology, Investigation. **Y Chen:** Investigation, Software. **D Cai**: Methodology, Investigation, Writing - review & editing. **Shouwen Chen**: Supervision, Writing - review & editing.

## Data availability statements

The datasets generated during and/or analyzed during the current study are available from the corresponding author on reasonable request.

## References

Abdel-Fattah WR, Chen Y, Eldakak A, Hulett FM (2005) *Bacillus subtilis* phosphorylated PhoP: direct activation of the E(sigma)^A^- and repression of the E(sigma)^E^-responsive *phoB*-P_S+V_ promoters during pho response. J Bacteriol, 187: 5166–5178.

Aggarwal S, Somani VK, Gupta V, Kaur J, Singh D, Grover A, Bhatnagar R (2017) Functional characterization of PhoPR two component system and its implication in regulating phosphate homeostasis in *Bacillus anthracis*. Biochim Biophys Acta Gen Subj, 1861: 2956–2970.

Allenby NE, O’Connor N, Pragai Z, Ward AC, Wipat A, Harwood CR (2005) Genome-wide transcriptional analysis of the phosphate starvation stimulon of *Bacillus subtilis*. J Bacteriol, 187: 8063–8080.

Cai D, Chen Y, He P, Wang S, Mo F, Li X, Wang Q, Nomura CT, Wen Z, Ma X et al (2018) Enhanced production of poly-gamma-glutamic acid by improving ATP supply in metabolically engineered *Bacillus licheniformis*. Biotechnol Bioeng, 115: 2541–2553.

Deng C, Wu Y, Lv X, Li J, Liu Y, Du G, Chen J, Liu L (2022) Refactoring transcription factors for metabolic engineering. Biotechnol Adv, 57: 107935.

Do TH, Suzuki Y, Abe N, Kaneko J, Itoh Y, Kimura K (2011) Mutations suppressing the loss of DegQ function in *Bacillus subtilis* (natto) poly-γ-glutamate synthesis. Appl Environ Microbiol, 77: 8249–8258.

Eldakak A, Hulett FM (2007) Cys303 in the histidine kinase PhoR is crucial for the phosphotransfer reaction in the PhoPR two-component system in *Bacillus subtilis*. J Bacteriol, 189: 410–421.

Freyre-González JA, Manjarrez-Casas AM, Merino E, Martinez-Nuñez M, Perez-Rueda E, Gutiérrez-Ríos R (2013) Lessons from the modular organization of the transcriptional regulatory network of *Bacillus subtilis*. BMC Syst Biol, 7.

Gaillard I, Slotboom DJ, Knol J, Lolkema JS, Konings WN (1996) Purification and reconstitution of the glutamate carrier GltT of the thermophilic bacterium *Bacillus stearothermophilus*. Biochemistry, 35: 6150–6156.

Han J, Gao X, Luo X, Zhu L, Zhang Y, Dong P (2023) The role of PhoP/PhoQ system in regulating stress adaptation response in Escherichia coli O157:H7. Food Microbiol, 112: 104244.

Hong LTT, Hachiya T, Hase S, Shiwa Y, Yoshikawa H, Sakakibara Y, Nguyen SLT, Kimura K (2019) Poly-γ-glutamic acid production of *Bacillus subtilis* (natto) in the absence of DegQ: A gain-of-function mutation in yabJ gene. Journal of Bioscience and Bioengineering, 128: 690–696.

Hu L, Zhao M, Hu W, Zhou M, Huang J, Huang X, Gao X, Luo Y, Li C, Liu K et al (2022a) Poly-γ-glutamic acid production by engineering a DegU quorum-sensing circuit in *Bacillus subtilis*. ACS Synth Biol.

Hu SY, He PH, Zhang YJ, Jiang M, Wang Q, Yang SH, Chen SW (2022b) Transcription factor DegU-mediated multi-pathway regulation on lichenysin biosynthesis in *Bacillus licheniformis*. Metab Eng, 74: 108–120.

Kobayashi K, Ogura M, Yamaguchi H, Yoshida K, Ogasawara N, Tanaka T, Fujita Y (2001) Comprehensive DNA microarray analysis of *Bacillus subtilis* two-component regulatory systems. J Bacteriol, 183: 7365–7370.

Li D, Hou L, Gao Y, Tian Z, Fan B, Wang F, Li S (2022) Recent advances in microbial synthesis of poly-gamma-glutamic acid: A review. Foods, 11.

Liu W, Hulett FM (1997) *Bacillus subtilis* PhoP binds to the *phoB* tandem promoter exclusively within the phosphate starvation-inducible promoter. J Bacteriol, 179: 6302–6310.

Liu W, Hulett FM (1998) Comparison of PhoP binding to the *tuaA* promoter with PhoP binding to other Pho-regulon promoters establishes a *Bacillus subtilis* Pho core binding site. Microbiology (Reading), 144: 1443–1450.

Martin JF, Rodriguez-Garcia A, Liras P (2017) The master regulator PhoP coordinates phosphate and nitrogen metabolism, respiration, cell differentiation and antibiotic biosynthesis: comparison in *Streptomyces coelicolor* and *Streptomyces avermitilis*. J Antibiot (Tokyo), 70: 534–541.

Martin JF, Sola-Landa A, Santos-Beneit F, Fernandez-Martinez LT, Prieto C, Rodriguez-Garcia A (2011) Cross-talk of global nutritional regulators in the control of primary and secondary metabolism in *Streptomyces*. Microb Biotechnol, 4: 165–174.

Mirouze N, Bidnenko E, Noirot P, Auger S (2015) Genome-wide mapping of TnrA-binding sites provides new insights into the TnrA regulon in *Bacillus subtilis*. Microbiology Open, 4: 423–435.

Nakano MM, Hoffmann T, Zhu Y, Jahn D (1998) Nitrogen and oxygen regulation of *Bacillus subtilis nasDEF* encoding NADH-dependent nitrite reductase by TnrA and ResDE. J Bacteriol, 180: 5344–5350.

Nakano MM, Zhu Y, Lacelle M, Zhang X, Hulett FM (2000) Interaction of ResD with regulatory regions of anaerobically induced genes in *Bacillus subtilis*. Mol Microbiol, 37: 1198–1207.

Nakano MM, Zuber P, Glaser P, Danchin A, Hulett FM (1996) Two-component regulatory proteins ResD-ResE are required for transcriptional activation of fnr upon oxygen limitation in *Bacillus subtilis*. J Bacteriol, 178: 3796–3802.

Ohsawa T, Tsukahara K, Ogura M (2009) *Bacillus subtilis* response regulator DegU is a direct activator of *pgsB* transcription Involved In *γ*-poly-glutamic acid synthesis. Biosci Biotechnol Biochem, 73: 2096–2102.

Pei JF, Li YX, Tang H, Wei W, Ye BC (2022) PhoP- and GlnR-mediated regulation of *metK* transcription and its impact upon S-adenosyl-methionine biosynthesis in *Saccharopolyspora erythraea*. Microb. Cell Fact., 21: 120–131.

Randazzo P, Aucouturier A, Delumeau O, Auger S (2017) Revisiting the in vivo GlnR-binding sites at the genome scale in *Bacillus subtilis*. BMC Res Notes, 10: 422.

Salzberg LI, Botella E, Hokamp K, Antelmann H, Maass S, Becher D, Noone D, Devine KM (2015) Genome-wide analysis of phosphorylated PhoP binding to chromosomal DNA reveals several novel features of the PhoPR-mediated phosphate limitation response in *Bacillus subtilis*. J Bacteriol, 197: 1492–506.

Santos-Beneit F (2015) The Pho regulon: a huge regulatory network in bacteria. Front Microbiol, 6.

Santos-Beneit F, Barriuso-Iglesias M, Fernandez-Martinez LT, Martinez-Castro M, Sola-Landa A, Rodriguez-Garcia A, Martin JF (2011) The RNA polymerase omega factor RpoZ is regulated by PhoP and has an important role in antibiotic biosynthesis and morphological differentiation in *Streptomyces coelicolor*. Appl Environ Microbiol, 77: 7586–7594.

Schau M, Eldakak A, Hulett FM (2004) Terminal oxidases are essential to bypass the requirement for ResD for full Pho induction in *Bacillus subtilis*. J Bacteriol, 186: 8424–8432.

Tran L-SP, Nagai T, Itoh Y (2000) Divergent structure of the ComQXPA quorum-sensing components: molecular basis of strain-specific communication mechanism in *Bacillus subtilis*. Mol Microbiol, 37: 1159–1171.

W Liu, Y Qi, Hulett FM (1998) Sites internal to the coding regions of *phoA* and *pstS* bind PhoP and are required for full promoter activity. Mol Microbiol, 28: 119–130.

Wei X, Zhou Y, Chen J, Cai D, Wang D, Qi G, Chen S (2015) Efficient expression of nattokinase in *Bacillus licheniformis*: host strain construction and signal peptide optimization. J Ind Microbiol Biotechnol, 42: 287–295.

Wicke D, Schulz LM, Lentes S, Scholz P, Poehlein A, Gibhardt J, Daniel R, Ischebeck T, Commichau FM (2019) Identification of the first glyphosate transporter by genomic adaptation. Environ Microbiol, 21: 1287–1305.

Xu Y, You D, Yao LL, Chu X, Ye BC (2019) Phosphate regulator PhoP directly and indirectly controls transcription of the erythromycin biosynthesis genes in *Saccharopolyspora erythraea*. Microb. Cell Fact., 18: 206.

Y Qi, Y Kobayashi, Hulett FM (1997) The *pst* operon of *Bacillus subtilis* has a phosphate-regulated promoter and is involved in phosphate transport but not in regulation of the Pho regulon. J Bacteriol, 179: 2534–2539.

Yuan H, Xu Y, Chen Y, Zhan Y, Wei X, Li L, Wang D, He P, Li S, Chen S (2019) Metabolomics analysis reveals global acetoin stress response of *Bacillus licheniformis*. Metabolomics : Official journal of the Metabolomic Society, 15: 25.

Zalieckas JM, Wray LV, Jr., Fisher SH (2006) Cross-regulation of the *Bacillus subtilis glnRA* and *tnrA* genes provides evidence for DNA binding site discrimination by GlnR and TnrA. J Bacteriol, 188: 2578–2585.

Zeng L, Choi SC, Danko CG, Siepel A, Stanhope MJ, Burne RA (2013) Gene regulation by CcpA and catabolite repression explored by RNA-Seq in Streptococcus mutans. PloS one, 8: e60465.

Zhan Y, Sheng B, Wang H, Shi J, Cai D, Yi L, Yang S, Wen Z, Ma X, Chen S (2018) Rewiring glycerol metabolism for enhanced production of poly-gamma-glutamic acid in *Bacillus licheniformis*. Biotechnology for biofuels, 11: 306.

Zhang Q, Chen Y, Gao L, Chen JG, Ma X, Cai D, Wang D, Chen S (2023) Enhanced production of poly-γ-glutamic acid via optimizing the expression cassette of *Vitreoscilla* hemoglobin in *Bacillus licheniformis*. Synth Syst Biotechnol, 7: 567–573.

Zhang Z, He P, Cai D, Chen S (2022) Genetic and metabolic engineering for poly-gamma-glutamic acid production: current progress, challenges, and prospects. World J Microbiol Biotechnol, 38: 208.

